# RAMP1 in Kupffer cells is a critical regulator in immune-mediated hepatitis

**DOI:** 10.1101/357582

**Authors:** Tomoyoshi Inoue, Yoshiya Ito, Nobuyuki Nishizawa, Koji Eshima, Ken Kojo, Fumisato Otaka, Tomohiro Betto, Sakiko Yamane, Kazutake Tsujikawa, Wasaburo Koizumi, Masataka Majima

## Abstract

The significanceF of the relationship between the nervous and immune systems with respect to disease course is increasingly apparent. Immune cells in the liver and spleen are responsible for the development of acute liver injury, yet the regulatory mechanisms of the interactions remain elusive. Calcitonin gene-related peptide (CGRP), which is released from the sensory nervous system, regulates innate immune activation via receptor activity-modifying protein 1 (RAMP1), a subunit of the CGRP receptor. Here, we show that RAMP1 in Kupffer cells (KCs) plays a critical role in the etiology of immune-mediated hepatitis. RAMP1-deficient mice with concanavalin A (ConA)-mediated hepatitis, characterized by severe liver injury accompanied by infiltration of immune cells and increased secretion of pro-inflammatory cytokines by KCs and splenic T cells, showed poor survival. Removing KCs ameliorated liver damage, while depleting T cells or splenectomy led to partial amelioration. Adoptive transfer of splenic T cells from RAMP1-deficient mice led to a modest increase in liver injury. Co-culture of KCs with splenic T cells led to increased cytokine expression by both cells in a RAMP1-dependent manner. Thus, immune-mediated hepatitis develops via crosstalk between immune cells. RAMP1 in KCs is a key regulator of immune responses.

## Introduction

The liver is exposed to high concentrations of products derived from the diet and gut microbiota, which in combination with bacterial endotoxins would normally trigger immune responses. Hepatic resident macrophages, called Kupffer cells (KCs), are the principal macrophages in the liver, where they reside in the sinusoids and are thus in a perfect position to monitor materials emanating from the intestine [1]. The liver is innervated by sympathetic, parasympathetic, and peptidergic nerves that contain both afferent and efferent nerve fibers. A number of neuropeptides are present in liver [2]. For example, neuronal fibers expressing substance P and calcitonin gene-related peptide (CGRP) are abundant in the tunica media of vessels in portal tracts, as well as in intra-lobular connective tissue [3]. The nervous system plays a major role in regulating immune homeostasis and inflammation, and the significanceF of the crosstalk between the nervous and immune systems with respect to injury outcome and/or disease course is increasingly apparent [4].

The immune system is tightly regulated by the nervous system, which releases several mediators (including neurotransmitters). CGRP, a 37 amino acid peptide, is produced in the neural body of dorsal root ganglion cells and released from sensory nerve endings [5]. CGRP binds to a specific receptor, a complex formed by receptor activity-modifying protein 1 (RAMP1) and calcitonin receptor-like receptor (CRLR) [6]. Binding of CGRP to its receptor triggers an increase in intracellular cyclic adenosine monophosphate levels, leading to vasodilation and transmission of pain signals [5,7]. Because CGRP receptors are expressed by immune cells, CGRP modulates immune responses via receptor binding [5]. Recent studies show that CGRP regulates the function of T cells [8], dendritic cells [9], and macrophages [10] during inflammation. CGRP suppresses production of tumor necrosis factor (TNF)α by lipopolysaccharide (LPS)-activated dendritic cells, thereby increasing production of interleukin (IL)-10 [9]. CGRP exerts similar immunosuppressive actions on T cells and macrophages in the colon [10]. These findings indicate that RAMP1 signaling and immune cells engage in crosstalk, thereby regulating immune responses.

In the liver, CGRP-deficient (*Calca*^−/−^) mice with immune-mediated hepatitis exhibit aggravated liver injury caused by concanavalin A (ConA), which enhances apoptosis mediated by interferon gamma (IFN γ) [11]. ConA hepatitis is mediated by activation of immune cells, including T lymphocytes [12] and KCs [13,14]. Because CGRP and its receptors are expressed in the liver [15] as well as spleen [16,17], RAMP1 signaling regulates ConA hepatitis via immune cells in the liver and spleen. However, it remains unknown about the contribution of RAMP1 signaling in the liver and spleen to the progression of immune-mediated hepatitis.

Here, we used RAMP1 knockout (*Ramp1^−/−^*) mice to explore the crosstalk between RAMP1 signaling and innate immunity in the liver and spleen during ConA-mediated hepatitis [9,18]. The results indicate that the initial liver injury is driven by interactions between splenic T cells and KCs, and that RAMP1 signaling in KCs orchestrates development of immune-mediated hepatitis. Thus, RAMP1 in KCs is a key modulator during development of immune-mediated hepatitis.

## Materials and methods

### Animals

Male C57Bl/6 WT mice were obtained from Clea Japan (Tokyo, Japan). Male *Ramp1^−/−^* mice were developed previously [9]. Mice were maintained at constant humidity (60 ± 5%) and temperature (25 ± 1°C) on a 12 h light/dark cycle. All animals were provided with food and water *ad libitum.* All experimental procedures were approved by the Animal Experimentation and Ethics Committee of the Kitasato University School of Medicine (2017-59), and were performed in accordance with the guidelines for animal experiments set down by the Kitasato University School of Medicine, which are in accordance with the “Guidelines for Proper Conduct of Animal Experiments” published by the Science Council of Japan. Mice used for survival studies were examined by animal care takers and the overall health status was checked by trained professionals. Mice were euthanized by pentobarbital sodium when they were found in a moribund state as identified by inability to maintain upright position and/or labored breathing. The mice for in vivo experiments were constantly watched throughout the experiment periods. Tissue collection procedures were performed under anesthesia with pentobarbital sodium. At the end of the experiments, the animals were euthanized by exsanguination under anesthesia with pentobarbital sodium followed by cervical dislocation.

### Animal procedures

Animals were fasted overnight and then intravenously (i.v.) injected (via the tail vein) with 20 mg/kg Con A (Merck KGaA, Darmstadt, Germany) dissolved in warm pyrogen-free saline (final concentration, 2.0 mg/ml) to induce hepatitis [18]. Mice were anaesthetized with pentobarbital sodium (60 mg/kg, intraperitoneally (i.p.)) at 0, 1, 3, 6, and 24 h after ConA administration (n = 6 for each time point), and blood was drawn. Levels of ALT were measured using a Dri-Chem 7000 Chemistry Analyzer System (Fujifilm, Tokyo, Japan). Immediately after blood collection, livers were excised and rinsed in saline. A small section of each liver was placed in 10% formaldehyde; the remaining liver was frozen in liquid nitrogen and stored at −80°C. For survival experiment, WT or *Ramp1^−/−^* mice were treated with ConA (n = 20 per group) and were monitored every 12 h up until 48 h after ConA administration. Some animals received a single subcutaneous injection of CGRP (5.0 µg/mouse in 200 µl of saline; Peptide Institute, Inc. Osaka, Japan) 30 min before treatment with Con A [15] or vehicle (saline).

#### Neutralization of TNFα and IFNγ

In another set of experiments, mice were injected i.p. with 100 μg of a neutralizing monoclonal antibody specific for mouse TNFα (eBioscience, San Diego, CA, USA) and 100 μg of a neutralizing monoclonal antibody specific for mouse IFNγ (eBioscience) 30 min before ConA administration.

#### Depletion of CD4+ T cells

Experimental animals were depleted of CD4+ cells using a rat anti-mouse CD4 monoclonal IgG2b antibody (clone GK1.5; BioLegend, San Diego, CA, USA). The antibody was administered i.p. (300 μg per mouse) 24 h before ConA administration. Control animals were treated with IgG isotype control antibodies (BioLegend).

#### Depletion of macrophages

Mice were injected i.v. with clodronate CL (200 μl/mouse; FormuMax Scientific, Inc., CA, USA) before 48 h ConA injection. Control groups were injected with control anionic liposomes (200 μl).

### Histology and immunohistochemistry

Excised liver tissues were fixed immediately with 10% formaldehyde prepared in 0.1 M sodium phosphate buffer (pH 7.4). Sections (3.5 µm thick) were prepared from paraffin-embedded tissues and either stained with hematoxylin and eosin (H&E) or immunostained with appropriate antibodies. Images of H&E-stained sections were captured under a microscope (Biozero BZ-7000 Series; KEYENCE, Osaka, Japan). Necrosis (expressed as a percentage of the total area) was estimated by measuring the necrotic area in the entire histological section using ImageJ software (US National Institutes of Health, Bethesda, MD, USA).

### Immunofluorescence analysis

Tissue samples were fixed with periodate-lysine-paraformaldehyde fixative at room temperature for 3 h. Following cryoprotection with 30% sucrose prepared in 0.1 M phosphate buffer (pH 7.2), sections (8 µm thick) were cut in a cryostat and incubated with Dako Protein Block Serum-Free solution (Glostrup, Denmark) at room temperature for 1 h to block non-specific binding. Sections were then incubated overnight at 4°C with a rabbit anti-mouse RAMP1 polyclonal antibody (Bioss Antibodies, Inc., Woburn, MA, USA), a rat anti-mouse Ly6C monoclonal antibody (Bio-Rad Laboratories, Inc., Puchheim, Germany), a rat anti-mouse CD4 monoclonal antibody (Bio-Rad Laboratories, Inc.), a rat anti-mouse CD3 monoclonal antibody (Bio-Rad Laboratories, Inc.), or a rat anti-mouse CD68 monoclonal antibody (Bio-Rad Laboratories, Inc.). After washing three times in PBS, the sections were incubated with a mixture of the following secondary antibodies for 1 h at room temperature: Alexa Fluor^®^ 488-conjugated donkey anti-rat IgG, Alexa Fluor^®^ 594-conjugated donkey anti-rabbit IgG, Alexa Fluor^®^ 488-conjugated donkey anti-goat IgG (all from Molecular Probes, OR, USA), and Alexa Fluor^®^ 594-conjugated goat anti-guinea pig IgG (Abcam plc, MA, USA). These antibodies were diluted in Antibody Diluent with Background-Reducing Components (Agilent, CA, USA). As a negative control, sections were incubated in Antibody Diluent with Background-Reducing Components in the absence of a primary antibody. Images were captured under a fluorescence microscope (Biozero BZ-9000 Series; KEYENCE). After labeling, six low-power optical fields (200× magnification) were randomly selected and the number of positive cells was counted. At least five animals were analyzed per marker.

### Real-time RT-PCR

Transcripts encoding *Ramp1*, *Calcrl*, *Calca*, *Tnf*, *Ifng*, *Ccl2*, *Ccl5*, *Ccl7*, and glyceraldehyde-3-phosphate dehydrogenase (*Gapdh*) were quantified by real-time RT-PCR analysis. Total RNA was extracted from mouse tissues and homogenized in TRIzol reagent (Life Technologies, Carlsbad, CA, USA). Single-stranded cDNA was generated from 1 μg of total RNA via reverse transcription using the ReverTra Ace^®^ qPCR RT Kit (TOYOBO Co., Ltd., Osaka, Japan), according to the manufacturer’s instructions. Quantitative PCR amplification was performed using SYBR Premix Ex Taq™ II (Tli RNaseH Plus; Takara Bio, Inc. Shiga, Japan). The gene-specific primers used for real-time RT-PCR were designed using Primer 3 software (http://primer3.sourceforge.net/) based on data from GenBank. The primers were as follows: 5’-CCATCTCTTCATGGTCACTGC-3’ (sense) and 5’-AGCGTCTTCCCAATAGTCTCC-3’ (antisense) for *Ramp1*; 5’-CTTCTGGATGCTCTGTGAAGG-3’ (sense) and 5’-CCCAGCCGAGAAAATAATACC-3’ (antisense) for *Calcrl*; 5’-TCTTCTCATTCCTGCTTGTGG-3’ (sense) and 5’-GATCTGAGTGTGAGGGTCTGG-3’ (antisense) for *Tnf*; 5’-ATCTGGAGGAACTGGCAAAAG-3’ (sense) and 5’-CGCTTATGTTGTTGCTGATGG-3’ (antisense) for *Ifng*; 5’-CGGAACCAAATGAGATCAGAA-3’ (sense) and 5’-TTGTGGAAAAGGTAGTGGATG-3’ (antisense) for *Ccl2*; 5’-CTGCTGCTTTGCCTACCTCTC-3’ (sense) and 5’-GTGACAAACACGACTGCAAGA-3’ (antisense) for *Ccl5*; 5’-TGCTTTCAGCATCCAAGTGTG-3’ (sense) and 5’-ACCGACTACTGGTGATCCTTC-3’ (antisense) for *Ccl7*; 5’-ACATCAAGAAGGTGGTGAAGC-3’ (sense) and 5’-AAGGTGGAAGAGTGGGAGTTG-3’ (antisense) for GAPDH. Data were normalized to GAPDH expression levels.

### Measurement of CGRP by ELISA

The concentrations of CGRP in liver and spleen tissues were measured with an ELISA kit (USCN Life Science Inc., Huston, TX, USA), according to the manufacturer’s instructions.

### Isolation of leukocytes from liver and spleen

Mice were anesthetized with pentobarbital sodium solution (60 mg/kg, i.p.), and the liver was perfused with perfusion buffer (10 ml, 1× Hank’s balanced salt solution) through the portal vein. Excised livers were placed immediately at room temperature in RPMI, minced into small pieces using scissors, and incubated in RPMI containing 0.05% collagenase (Type IV; Sigma Chemical Co., St. Louis, MO, USA) at 37°C for 20 min. The tissue was then pressed through a 70 μm cell strainer. The cells were centrifuged at 2600 rpm for 10 min at 4°C, and pelleted cells were resuspended in PBS. Hepatic leukocytes were isolated from liver homogenates by density-gradient centrifugation on 33% Percoll™ (GE Healthcare Life Sciences, Piscataway, NJ, USA), as previously reported [19,20]. Non-parenchymal cells were collected from the interface between the 33% and 66% Percoll™ density cushions and centrifuged at 2700 rpm for 30 min at 4°C. The spleen was also collected and placed immediately in ice-cold RPMI. The tissue was pressed through a 70 μm cell strainer, and erythrocytes were disrupted in lysis buffer. Viable, nucleated cells were counted by trypan blue exclusion and diluted to a uniform cell density.

### Cell culture and co-culture conditions

F4/80+ cells were isolated from the liver and spleen using the mouse F4/80+ Isolation Kit (Miltenyi Biotec, Auburn, CA, USA), according to the manufacturer’s instructions. Then, F4/80+ cells were cultured in RPMI 1640 medium containing 10% fetal calf serum and plated in 6-well plates (1.0 × 10^6^ cells per well). After 24 h, cells were stimulated for 3 h with ConA (2 μg/ml) with or without CGRP (1 or 10 nM) (Peptide Institute) in RPMI 1640 medium. CD4+ cells were isolated from the spleen using the mouse CD4+ T cell Isolation Kit (Miltenyi Biotec), according to the manufacturer’s instructions. Isolated CD4+ cells (10^6^ cells/ml) were stimulated with 1 mg/ml anti-CD3 (BioLegend) and 1 mg/ml anti-CD28 (BioLegend). After 24 h, cells were stimulated for 1 h with ConA with or without CGRP (Peptide Institute) in RPMI 1640 medium. For the co-culture experiments, isolated KCs from liver and/or isolated CD4+ T cells from spleen were seeded into plates. The co-culture was maintained for 24 h, after which co-cultured cells were incubated for 1 h with or without ConA (2 µg/ml) in the presence/absence of CGRP (1 or 10 nM) in advanced RPMI 1640 (Thermo Fisher Scientific, Inc. Waltham, MA, USA). The F4/80+ cells and CD4+ cells were then harvested and homogenized in TRIzol (Life Technologies), and mRNA levels were measured by real-time RT-PCR.

### Flow cytometry analysis

Cells were incubated with the 2.4G2 mAb (anti-cγRIII/II) to block non-specific binding of the primary mAb. Then, cells were stained with a combination of the following fluorochrome-conjugated antibodies: anti-CD45 (clone 30-F11, BioLegend), anti-Ly6G (clone 1A8, BioLegend), anti-Ly6C (clone HK 1.4, BioLegend), anti-CD11b (clone M1/70, BioLegend), anti-F4/80 (clone A3-1, Bio-Rad), anti-CD3 (clone 17A2, BioLegend), anti-CD4 (clone GK1.5, BioLegend), and anti-CD8 (clone 53-6.7, BioLegend). Tubes were placed in the dark on ice for 30 min. Pellets were washed twice with PBS. For flow cytometric analysis, cells were initially gated on forward-scatter (FSC) and side-scatter (SSC) and then gated on CD45+ cells. Cells positive for 7-aminoactinomycin D (7-AAD; BioLegend) were electronically gated from the analysis as dead cells. Samples were measured on a FACSVerse™ (BD, Franklin Lakes, NJ, USA). The data were analyzed using Kaluza software v1.3 (Beckman Coulter, Brea, CA, USA) [19,20]. For intracellular cytokine staining, cells were incubated for 4 h at room temperature with Golgi Plug (BD Biosciences, San Jose, CA, USA) to block intracellular protein transport. Then, cells were incubated with the 2.4G2 mAb. Saturating concentrations of APC-labeled anti-F4/80 (clone A3-1, Bio-Rad) and APC/Cy7-labeled anti-CD4 (clone GK1.5, BioLegend) antibodies were added. Following washing, cells were fixed and permeabilized prior to intracellular staining of cytokines. After fixation and permeabilization, cells were incubated with PE-labeled anti-TNFα (BioLegend) and FITC-labeled anti-IFNγ (BioLegend) antibodies.

### Adoptive transfer of T cells

Isolated splenic CD4-positive cells (T cells) from WT and *Ramp1^−/−^* mice were injected into WT mice (i.v. at a dose of 5 × 10^6^ cells/200 μ1, of PBS). Seven days later, recipient mice were treated with ConA. At 24 h post-treatment, serum and liver samples were collected for analyses.

### Splenectomy

For splenectomy, the abdominal wall of mice was opened through a left subcostal minimal incision under ether anesthesia. The splenic arteries and veins were ligated at the splenic hilum using 3–0 silk and then divided. The resected spleen was removed, and the abdominal incision was closed. All surgical procedures were performed under sterilized conditions. For the sham operation, the abdominal wall was similarly opened and then closed immediately after identifying the spleen.

### Statistical analysis

All results are expressed as the mean ± standard deviation (SD). All statistical analyses were performed using GraphPad Prism software, version 6.07 (GraphPad Software, La Jolla, CA, USA). Unpaired two-tailed Student’s t-test was used to compare data between two groups, and one-way analysis of variance, followed by Bonferroni’s post-hoc test, was used to compare data between multiple groups. The survival rates of WT and *Ramp1^−/−^* mice were compared using Kaplan-Meier survival analysis and log-rank tests. A P-value < 0.05 was considered statistically significant.

## Results

### ConA-mediated hepatitis is exacerbated in *Ramp1^−/−^* mice

To elucidate the functional role of RAMP1 signaling during immune-mediated liver injury, we treated WT and *Ramp1^−/−^* mice with ConA. First, we examined the impact of RAMP1 deficiency on ConA-induced mortality by monitoring survival every 12 h up until 48 h after ConA administration. We found that 40% of *Ramp1*^−/−^ mice died within 24 h, whereas all of the WT mice survived (Fig 1A). These results suggest that *Ramp1^−/−^* mice are highly susceptible to acute liver injury caused by ConA. Based on these results, we compared WT and *Ramp1^−/−^* mice within 24 h of ConA treatment in all subsequent experiments.

**Fig 1.**
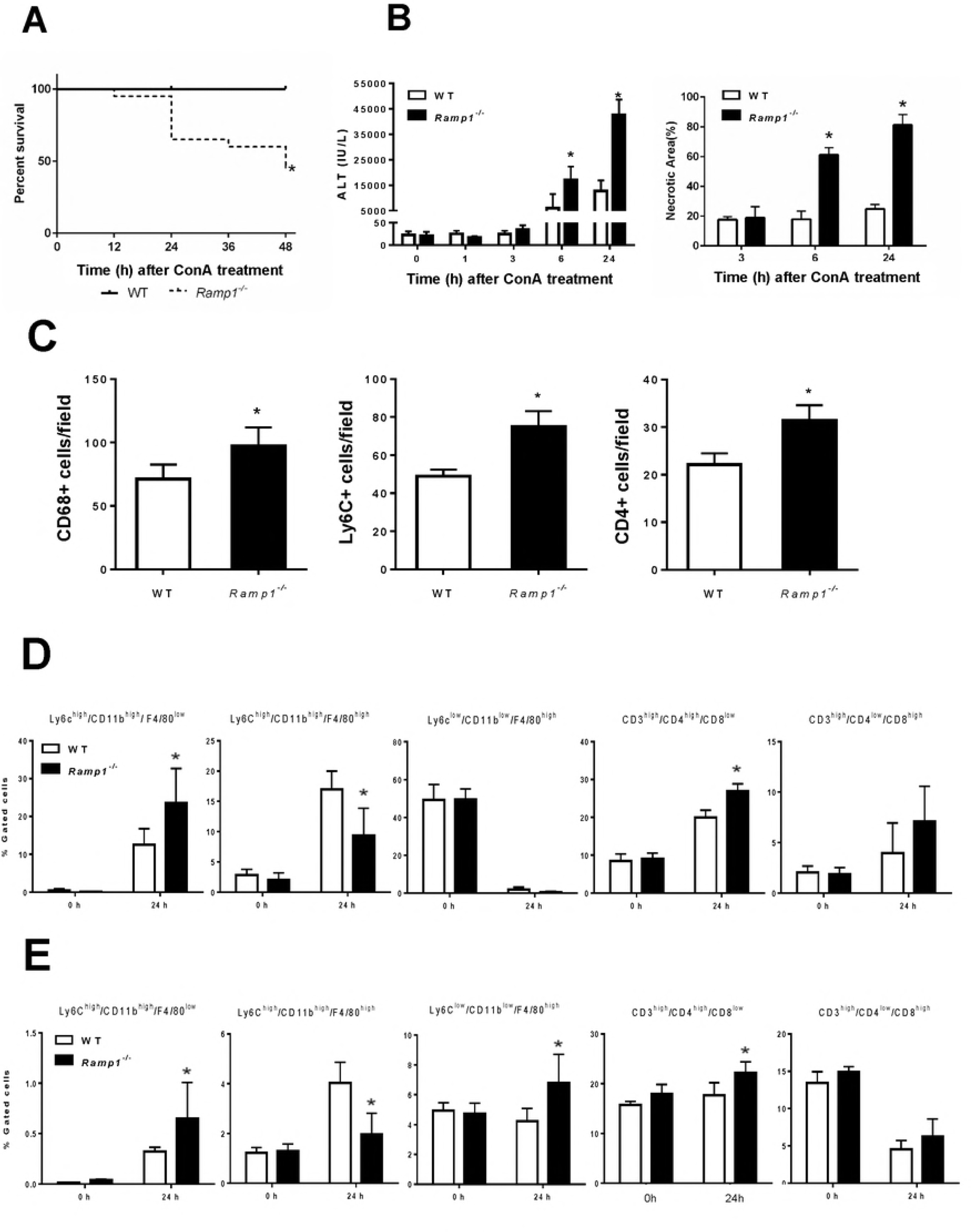
Deficient RAMP1 signaling exacerbates ConA-induced liver injury via infiltration of inflammatory cells into the liver. *Ramp1*^−/−^ and WT mice were administered ConA (20 mg/kg) by intraperitoneal (i.p.) injection. (a) Kaplan-Meier survival curves for *Ramp1^−/−^* mice (n = 20) and WT mice (n = 20) at various time points after receiving ConA. Survival data analyzed by log-rank test *p < 0.05 *vs.* WT mice. (b) Serum levels of ALT and the area of hepatic necrosis (expressed as a percentage) after ConA administration. Data are expressed as the mean ± SD (n = 6 mice per group). *p < 0.05 *vs.* WT mice. Serum samples at 0, 1, 3, 6, and 24 h after ConA administration were collected, and ALT levels were measured. (c) Numbers of hepatic Ly6C+ cells, CD68+ cells, and CD4+ cells at 24 h after ConA administration. Data are expressed as the mean ± SD of six mice per group. *p < 0.05 *vs*. WT mice. (d, e) Flow cytometry analysis of the percentage of Ly6C^high^/CD11b^high^/F4/80^low^ cells, Ly6C^high^/CD11b^high^/F4/80^high^ cells Ly6C^low^/CD11b^low^/F4/80^high^ cells, CD3^high^/CD4^high^/CD8^low^ cells, and CD3^high^/CD4^low^/CD8^high^ cells within the CD45+ cell population in the liver (d) and spleen (e) at 0 and 24 h after ConA treatment. Data are expressed as the mean ± SD of six mice per group. *p < 0.05 *vs.* WT mice.

Next, we examined the role of RAMP1 signaling in ConA-induced hepatitis by biochemical and histological evaluation of liver injury. Within 6 h of ConA treatment, ALT levels in WT mice began to increase and were elevated markedly at 24 h post-treatment (Fig 1B). By contrast, liver injury in *Ramp1^−/−^* mice was more severe than that in WT mice, as reflected by markedly higher ALT levels. The same was true for hepatic necrosis, as indicated by the larger necrotic areas in the liver of *Ramp1^−/−^* mice (Fig 1B and S1 Fig).

Because inflammatory cells, including T cells and macrophages, are responsible for development of ConA hepatitis via production of pro-inflammatory cytokines, we examined accumulation of inflammatory cells in the liver and spleen. Immunofluorescence analysis revealed that, at 24 h after ConA treatment, accumulation of inflammatory cells (including Ly6C+, CD68+, and CD4+ cells) in *Ramp1^−/−^*-livers was much higher than that in WT livers (Fig 1C and S1 Fig). During the early phase of ConA hepatitis, the number of hepatic CD3+ cells increased minimally (S1 Fig). Next, we examined the recruited inflammatory cells using flow cytometry analysis (Fig 1D 1E, S2 Fig). Exposure to ConA for 24 h caused hepatic accumulation of monocytes (defined as Ly6C^high^/CD11b^high^/F4/80^low^) or recruited macrophages (Ly6C^high^/CD11b^high^/F4/80^high^) in the livers of both genotypes. There were more Ly6C^high^/CD11b^high^/F4/80^low^ cells in *Ramp1^−/−^* mice than in WT mice, but fewer Ly6C^high^/CD11b^high^/F4/80^high^ cells (Fig 1D). These observations were associated with higher hepatic levels of chemokines, including C-C motif chemokine ligand 2 (*Ccl2*), *Ccl5*, and *Ccl7*, in *Ramp1^−/−^* mice than in WT mice (S3 Fig). By contrast, KCs (defined as Ly6C^low^/CD11b^low^/F4/80^high^) were markedly reduced in both genotypes (Fig 1D). There were more CD4+ T cells (CD3^high^/CD4^high^/CD8^low^) in the liver of *Ramp1^−/−^* mice than in that of WT mice (Fig. 1D). However, there were no significant differences between the genotypes in terms of the number of CD8+ T cells (CD3^high^/CD4^low^/CD8^high^) in the liver (Fig 1D). These results suggest that RAMP1 protects the liver from ConA hepatotoxicity by inhibiting accumulation of inflammatory cells.

There were more monocytes (Ly6C^high^/CD11b^high^/F4/80^low^) in the spleen of *Ramp1^−/−^* mice than in WT mice, but fewer macrophages (Ly6C^high^/CD11b^high^/F4/80^high^) (Fig 1E). However, the percentage of these cells was small in both genotypes. There was a very small increase in the number of resident macrophages (Ly6C^low^/CD11b^low^/F4/80^high^) in *Ramp1^−/−^* mice, along with moderate increases in the number of CD4+ T cells. By contrast, the number of CD8+ T cells fell in both genotypes (Fig 1E).

### Expression of CGRP and RAMP1 during ConA-induced hepatitis

Next, we examined expression of RAMP1 in the liver and spleen from WT mice to investigate the potential role of RAMP1 in ConA-induced liver injury. Real-time PCR analysis revealed that the amount of *Ramp1* mRNA in the liver increased transiently at 1 h after ConA administration before declining thereafter, reaching a nadir at 6 h, and recovering to baseline levels at 24 h (S4 Fig). A gradual downregulation of *Ramp1* expression over time was observed in the spleen after ConA treatment (S4 Fig). A similar trend was observed for *Calcrl*, another subunit of the CGRP receptor, in the liver and spleen after ConA administration S4 Fig); however, the levels of *Calcrl* did not differ between *Ramp1^−/−^* and WT mice. Next, we tried to identify the cellular sources of RAMP1. CGRP, a ligand of RAMP1, was also identified in nerve fibers within the liver and spleen (S4 Fig). CGRP levels fell over time after ConA treatment, and hepatic levels at 3, 6, and 24 h in *Ramp1^−/−^* mice were lower than those in WT mice (S4 Fig). CGRP expression in the spleen of both genotypes dropped at 1 h, before gradually returning to normal levels (S4 Fig); however, there were no significant differences between the genotypes. These results suggest that downregulation of CGRP/RAMP1 is associated with ConA hepatitis, and that RAMP1 in macrophages and T cells might play an important role in this process.

### Increased expression of pro-inflammatory cytokines in *Ramp1*^−/−^ mice with ConA hepatitis

Because pro-inflammatory cytokines, including TNFα and IFNγ, contribute to ConA hepatitis, we next measured *Tnf* and *Ifng* mRNA in the liver and spleen of mice with ConA hepatitis. Real-time PCR analysis revealed that WT and *Ramp1^−/−^* mice showed maximum expression of *Tnf* and *Ifng* at 1 h after ConA administration (Fig 2A). *Ramp1^−/−^* mice showed higher expression of *Tnf* and *Ifng* in the liver than WT mice. After 1 h, the levels of both cytokines fell. *Tnf* and *Ifng* mRNA expression was also higher in the spleen of *Ramp1^−/−^* mice (Fig 2B) than in that of WT mice, indicating that *Ramp1* signaling regulates expression of cytokines in the spleen, thereby contributing to ConA hepatitis.

**Fig 2.**
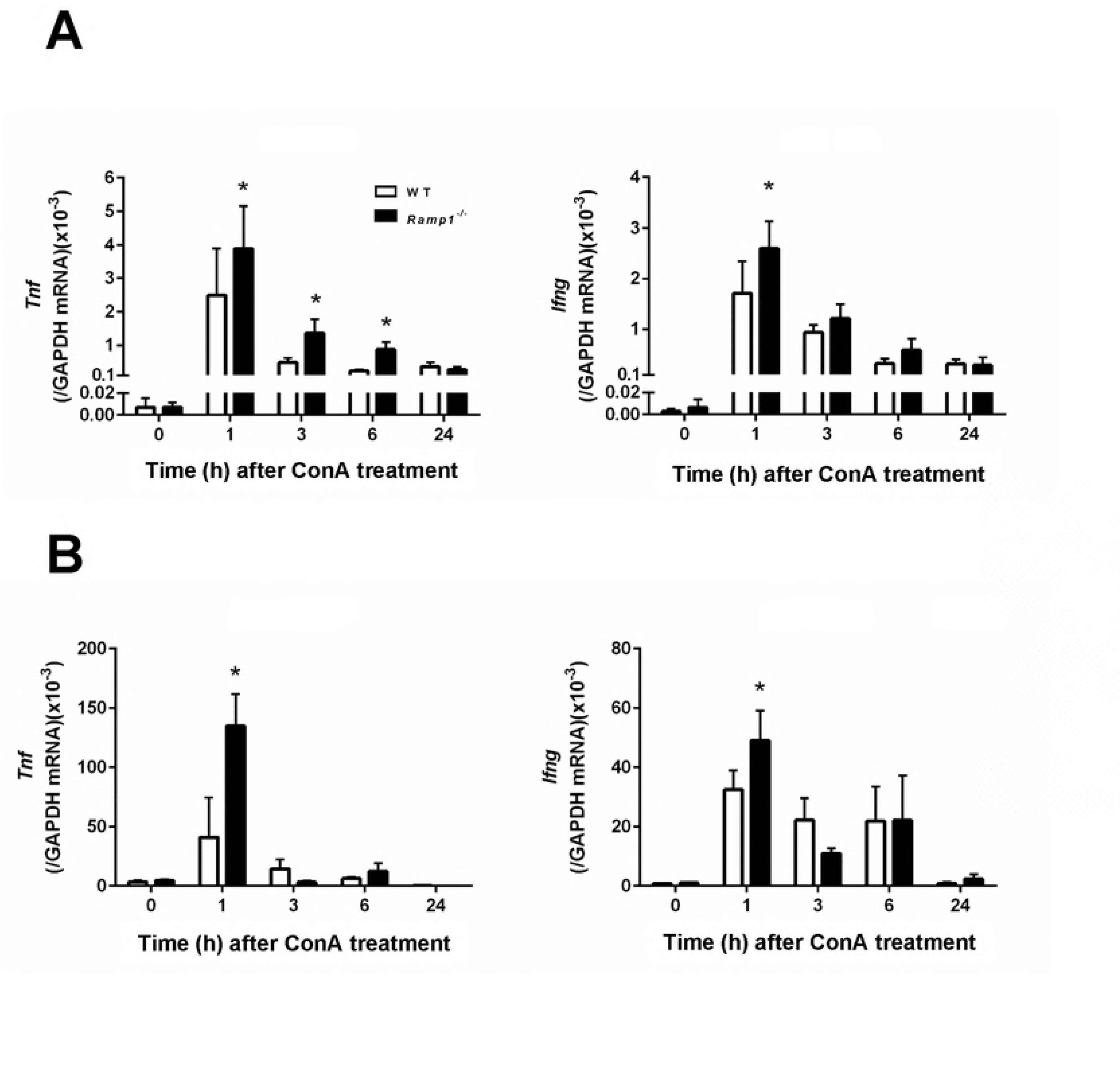
Increased expression of pro-inflammatory cytokines in *Ramp1^−/−^* mice with ConA hepatitis. Time course of changes in mRNA levels of *Tnf* and *Ifng* in the liver (a) and spleen (b) of WT and *Ramp1^−/−^* mice after ConA treatment. Data are expressed as the mean ± SD from six mice per group. *p < 0.05 *vs.* WT mice.

### Splenocytes play a role in ConA hepatitis

To investigate the possible role of splenocytes in development of ConA hepatitis, mice were subjected to splenectomy or a sham operation immediately prior to administration of ConA. At 6 h post-administration, the levels of ALT in splenectomized WT and *Ramp1^−/−^* mice decreased by 95% and 80%, respectively (Fig 3A). Thus, splenectomy ameliorated liver injury (as measured by ALT levels) in WT and *Ramp1^−/−^* mice. The mRNA levels of *Tnf* at 1 h after ConA treatment were decreased in splenectomized *Ramp1^−/−^* mice, but not splenectomized WT mice (Fig 3B). The mRNA levels of *Ifng* were downregulated in splenectomized WT and *Ramp1^−/−^* mice (Fig 3B). These results indicate that splenectomy suppresses the early phase of ConA hepatitis and that the spleen is the source of cytokines during ConA hepatitis.

**Fig 3.**
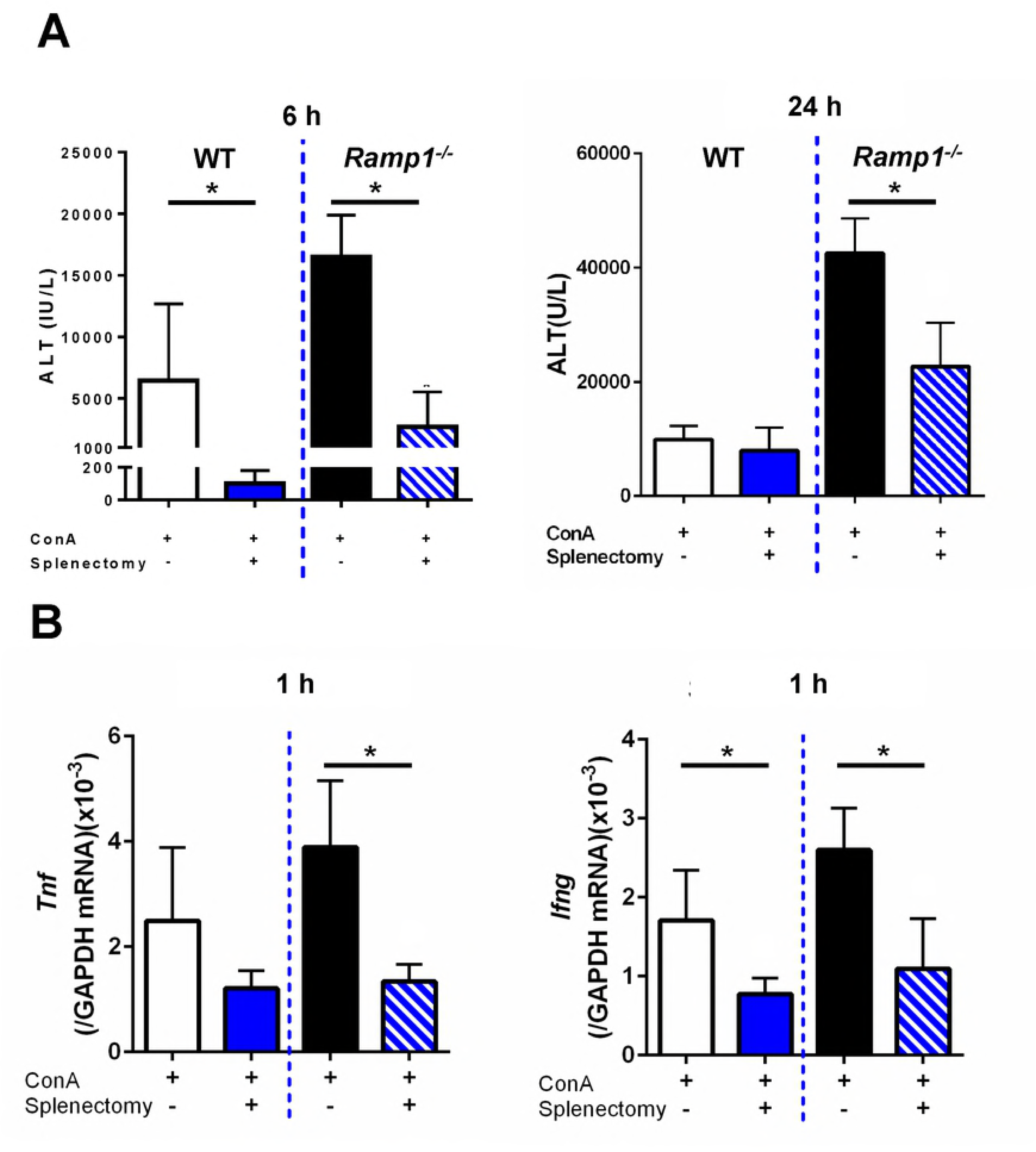
Effects of splenectomy on ALT and pro-inflammatory cytokines levels in mice with ConA hepatitis. (a) Splenectomy reduced ALT levels in WT and *Ramp1^−/−^* mice at 6 h (left) and 24 h (right) after ConA treatment. ConA was administered immediately after splenectomy. Data are expressed as the mean ± SD from 4–6 mice per group. *p < 0.05. (b) Effect of splenectomy on *Tnf* and *Ifng* mRNA levels in liver from WT and *Ramp1^−/−^* mice at 1 h after ConA administration. ConA was administered immediately after splenectomy. Data are expressed as the mean ± SD from 4–6 mice per group. *p < 0.05.

To better understand which cells are responsible for producing pro-inflammatory cytokines in the spleen, we measured cytokine production by splenic macrophages and T cells. Flow cytometry analysis revealed that the numbers of TNFα- and IFNγ-producing F4/80+ macrophages in *Ramp1^−/−^* mice at 1 h post-ConA administration were higher than in WT mice (Fig 4A). The numbers of TNFα- and IFNγ-producing CD4+ T cells in *Ramp1^−/−^* mice were also higher than those in WT mice (Fig 4B). These results indicate that RAMP1 signaling in splenic macrophages and T cells inhibits generation of TNFα and IFNγ in response to ConA. In addition, more CD4+ T cells than macrophages expressed these cytokines, suggesting that CD4+ T cells are the main source of TNFα and IFNγ during ConA hepatitis.

**Fig 4.**
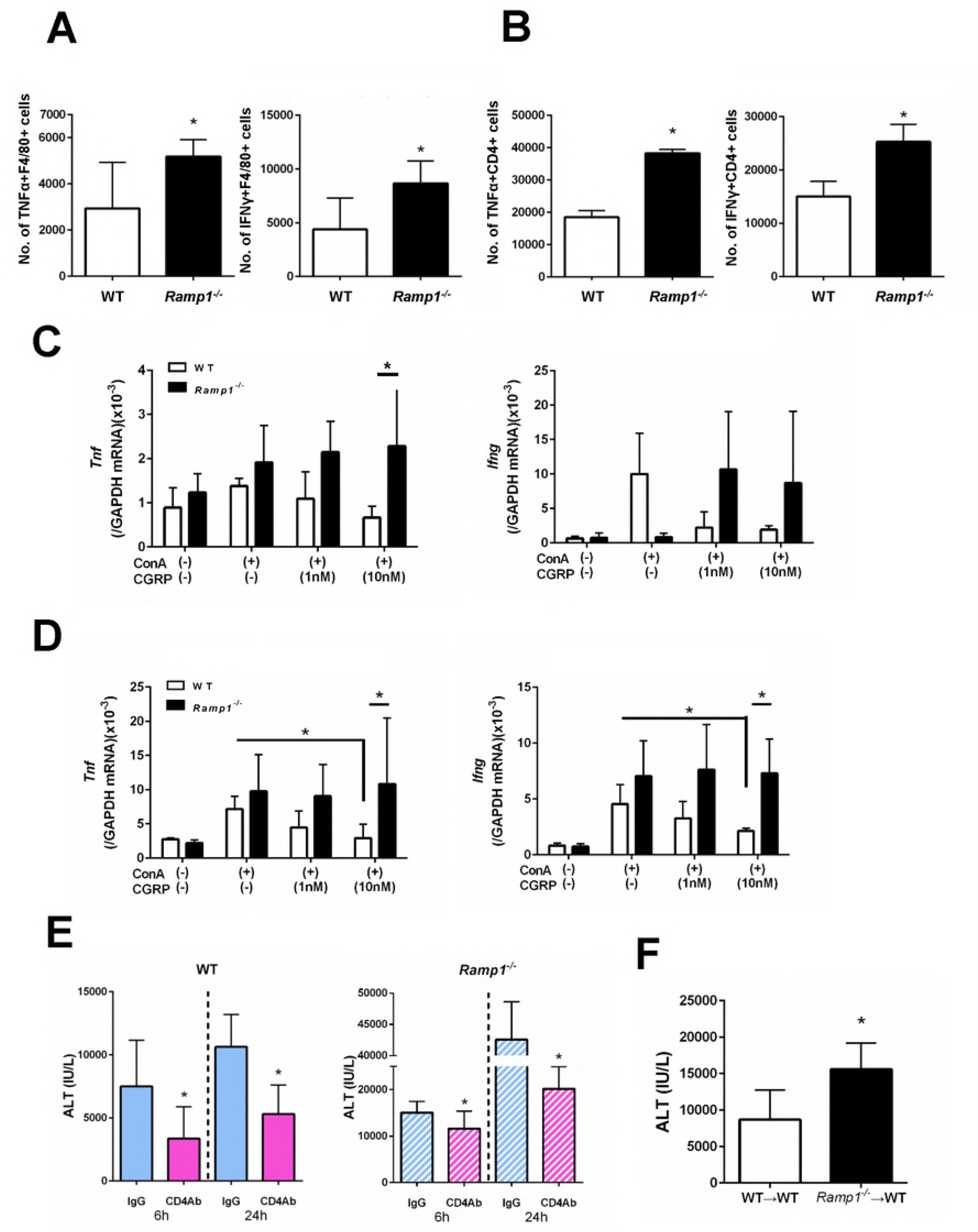
RAMP1 signaling regulates expression of pro-inflammatory cytokines by splenic macrophages and T cells. (a, b) The numbers of TNFα- and IFNγ-expressing cells within splenic F4/80+ cells (a) and splenic CD4+ T cells (b) populations from WT and *Ramp1^−/−^* mice at 1 h after ConA treatment. Expression of TNFα and IFNγ was analyzed by flow cytometry. Data are expressed as the mean ± SD from six mice per group. *p < 0.05 *vs.* WT mice. (c) Amounts of *Tnf* and *Ifng* mRNA expressed by splenic F4/80+ cells isolated from WT and *Ramp1^−/−^* mice. Isolated splenic macrophages were stimulated with CGRP (1 and 10 nM) with or without ConA for 3 h. Macrophages were isolated by magnetic-activated cell sorting (MACS). Data are expressed as the mean ± SD from 4–5 mice per group. *p < 0.05. (d) Expression of *Tnf* and *Ifng* mRNA by splenic CD4+ T cells isolated from WT and *Ramp1^−/−^* mice. Isolated splenic CD4+ T cells were stimulated with CGRP with or without ConA for 1 h. CD4+ T cells were isolated by magnetic-activated cell sorting (MACS). Data are expressed as the mean ± SD from 4–5 mice per group. *p < 0.05. (e) Serum ALT levels after deletion of CD4+ T cells from WT and *Ramp1^−/−^* mice during ConA hepatitis. Anti-CD4 antibodies or isotype IgG was administered 24 h before ConA treatment. Data are expressed as the mean ± SD from 3–6 mice per group. *p < 0.05 *vs.* IgG-treated mice. (f) Adoptive transfer of CD4+ T cells from *Ramp1^−/−^* mice into WT mice (*Ramp1^−/−^*→WT). Data are expressed as the mean ± SD from 3–6 mice per group. *p < 0.05 *vs.* WT→WT.

To better understand the role of RAMP1 signaling in splenic macrophages and CD4+ T cells, we isolated these cells from WT and *Ramp1^−/−^* mice and cultured them. Upon stimulation with ConA, CGRP downregulated *Tnf* and *Ifng* in splenic macrophages from WT mice, but upregulated them in splenic macrophages from *Ramp1^−/−^* mice (Fig 4C). Similarly, levels of *Tnf* and *Ifng* in splenic CD4+ T cells from WT mice were reduced, while those in splenic T cells from *Ramp1^−/−^* mice increased (Fig 4D). These results suggest that RAMP1 signaling in splenic macrophages and T cells downregulates expression of mRNA encoding *Tnf* and *Ifng*.

Splenectomy failed to reduce ALT levels in WT mice at 24 h after administration of ConA. By contrast, splenectomy decreased ALT levels in *Ramp1^−/−^* mice by 50% (see Fig 3A). At 6 h after ConA administration, splenectomy protected against ConA hepatitis in WT mice during the early phase, but not the late phase, of injury. Regarding RAMP1 signaling, splenectomy led to a partial improvement in ConA hepatitis in *Ramp1^−/−^* mice, suggesting that RAMP1 signaling in splenocytes participates in ConA-mediated hepatitis. Taken together, the results suggest that RAMP1-expressing cells other than splenocytes are required for amplification of liver injury elicited by ConA.

To further clarify the role of T cells (CD4+ cells) in ConA hepatitis, we treated WT and *Ramp1^−/−^* mice with an anti-CD4 antibody 24 h before ConA treatment. The anti-CD4 antibody suppressed ConA liver injury in WT and *Ramp1^−/−^* mice at 6 and 24 h post-ConA administration. The anti-CD4 antibody caused a partial reduction in ALT levels, suggesting that RAMP1 signaling in CD4+ T cells makes a limited contribution to ConA hepatitis (Fig 4E). These results were consistent with those observed in mice subjected to splenectomy.

To elucidate the contribution of RAMP1 signaling in splenic T cells to ConA hepatitis, we isolated splenic CD4+ T cells from *Ramp1^−/−^* mice and adoptively transferred them into WT mice. Transfer of *Ramp1*-deficient CD4+ T cells increased ALT levels (Fig 4F). However, the magnitude of liver injury in WT mice receiving *Ramp1*-deficient CD4+ T cells was lower than that in Ramp1 null mice, suggesting that Ramp1-deficient CD4+ T cells play a role (at least partially) in exacerbating liver injury in global knockout *Ramp1* mice. Conversely, *Ramp1*-deficient cells other than splenic CD4+ T cells exacerbate ConA hepatitis in *Ramp1^−/−^* mice.

### RAMP1 in KCs protects the liver from ConA hepatitis

Next, we examined whether hepatic macrophages aggravate ConA hepatitis in *Ramp1^−/−^* mice. In agreement with previous studies, we found that pre-treatment with clodronate liposomes (CL) attenuated ConA hepatitis in WT and *Ramp1^−/−^* mice (Fig 5A and 5B), suggesting that KCs are involved in induction and progression of ConA hepatitis. Treatment with CL also reduced *Tnf* and *Ifng* expression in WT (Fig 5C and 5E) and *Ramp1^−/−^* (Fig 5D and 5F) mice. At 24 h post-ConA treatment, CL suppressed accumulation of inflammatory cells (CD68+ and Ly6C+ cells) in WT and *Ramp1^−/−^* mice (S5 Fig), which was associated with downregulation of chemokines (*Ccl2* and Ccl7) (S5 Fig). These results suggest that RAMP1 signaling in KCs is responsible for ConA-mediated hepatitis.

**Fig 5.**
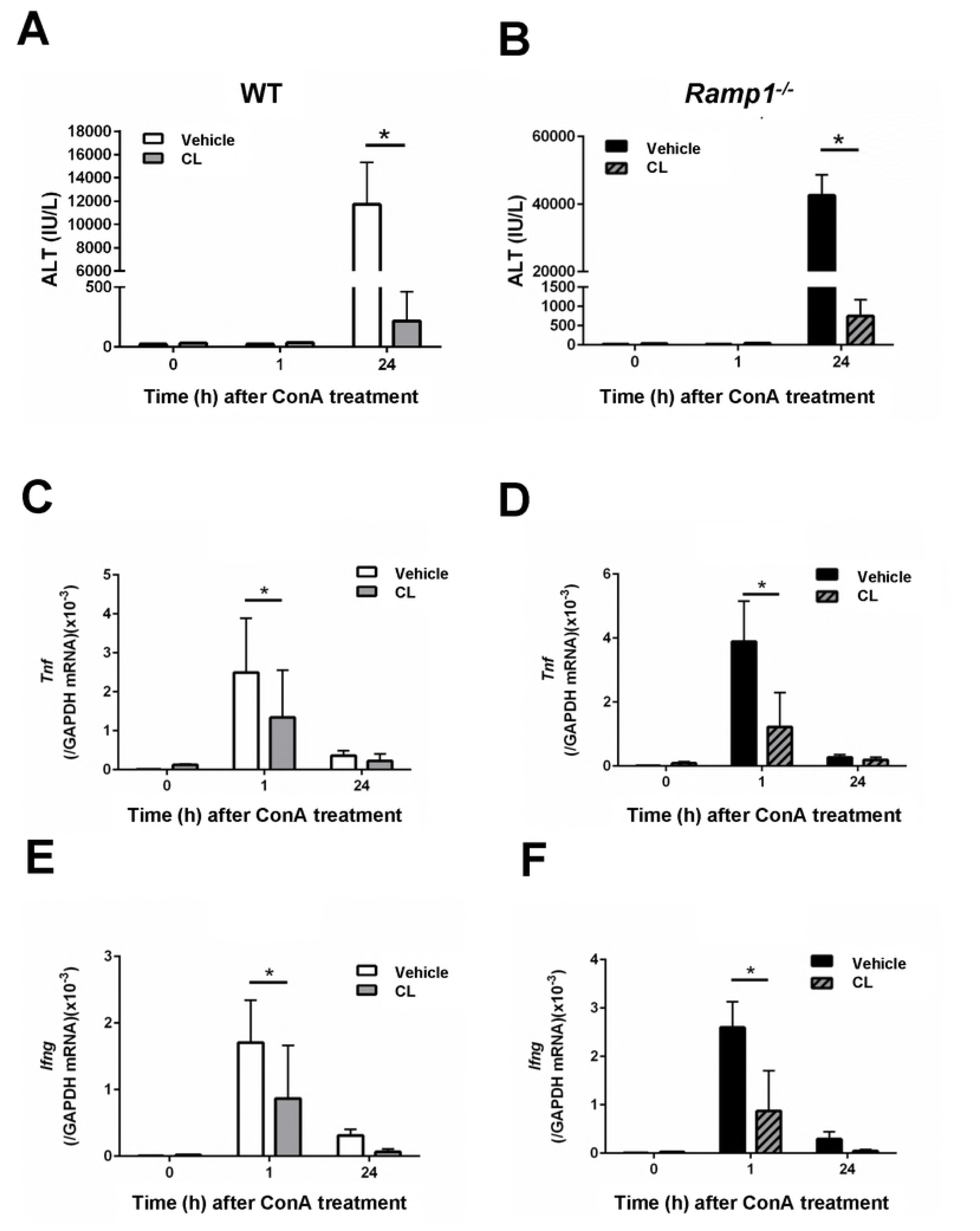
Deleting macrophages attenuates ConA-induced liver injury and decreases production of pro-inflammatory cytokines. (a, b) ALT levels in WT mice (a) and *Ramp1^−/−^* mice (b) treated with clodronate liposomes (CL). Mice were treated with CL or vehicle (PBS) 48 h before ConA administration. Data are expressed as the mean ± SD from 3–6 mice per group. *p < 0.05. (c–f) Amounts of *Tnf* mRNA in WT mice (c) and *Ramp1^−/−^* mice (d) treated with CL, and amounts of Ifng mRNA in WT mice (e) and *Ramp1^−/−^* mice (f) treated with CL. Data are expressed as the mean ± SD from 3–6 mice per group. *p < 0.05.

TNFα and IFNγ are key causative agents of ConA hepatitis. To elucidate the role of TNFα/IFNγ in our model, WT mice were treated with neutralizing anti-TNFα and anti-IFNγ antibodies prior to ConA administration. Blocking TNFα/IFNγ prevented ConA-induced liver injury, as evidenced by reduced ALT levels, less hepatic necrosis, and reduced infiltration of the liver by monocytes/macrophages and T cells (S6 Fig). These results indicate that TNFα/IFNγ play a critical role in mediating ConA-induced hepatitis.

Next, we measured TNFα and IFNγ secreted by hepatic macrophages and T cells using flow cytometry analysis. At 1 h after ConA treatment, the number of TNFα-expressing KCs in *Ramp1^−/−^* mice was higher than that in WT mice (Fig 6A). By contrast, there was no significant difference in the number of TNFα-expressing CD4+ T cells between the genotypes. The number of IFNγ-expressing KCs in *Ramp1^−/−^* mice was also greater than that in WT mice (Fig 6B). However, no significant difference in the number of IFNγ-expressing CD4+ T cells between the genotypes was observed. These results suggest that, in the liver, RAMP1 signaling downregulates cytokine production by KCs.

**Fig 6.**
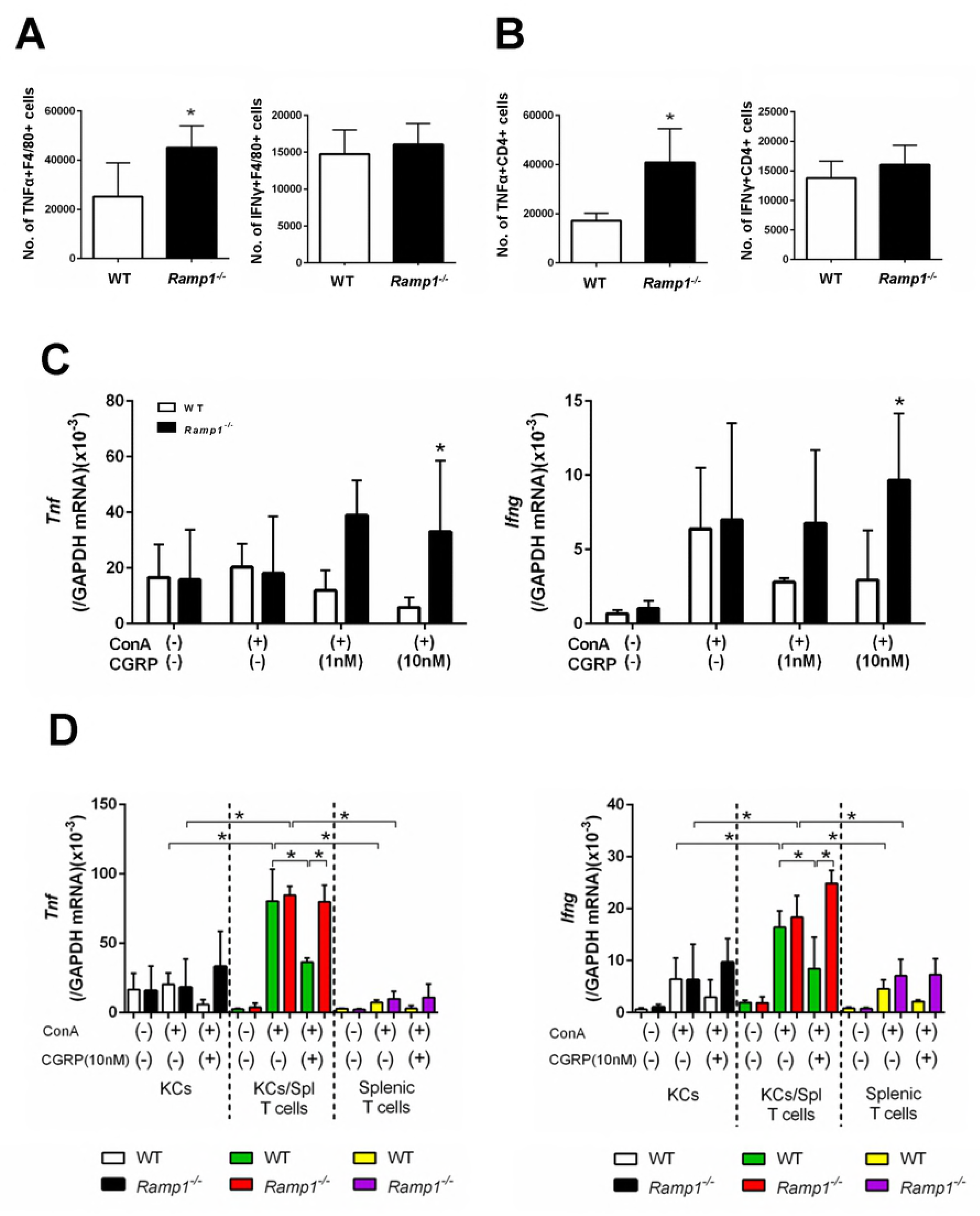
RAMP1 signaling regulates expression of pro-inflammatory cytokines in hepatic macrophages and splenic T cells. (a, b) The numbers of TNFα- and IFNγ-producing F4/80+ cells (a) and the numbers of TNFα- and IFNγ producing CD4+ T cells (b) in the liver from WT and *Ramp1^−/−^* mice at 1 h after ConA treatment. Expression of TNFα and IFNγ was analyzed by flow cytometry. Data are expressed as the mean ± SD from six mice per group. *p < 0.05 vs. WT mice. (c) Amounts of *Tnf* and *Ifng* mRNA in KCs isolated from WT and *Ramp1^−/−^* mice. Isolated KCs were stimulated with CGRP (1 and 10 nM) with or without ConA for 3 h. KCs were isolated by magnetic-activated cell sorting (MACS). Data are expressed as the mean ± SD from 4–5 mice per group. *p < 0.05 *vs.* WT mice. (d) Amounts of Tnf and Ifng mRNA in KCs and/or splenic CD4+ T cells isolated from WT and *Ramp1^−/−^* mice. KCs co-cultured with splenic CD4+ T cells were stimulated with CGRP with or without ConA for 1 h. Data are expressed as the mean ± SD from 4–5 mice per group. *p < 0.05.

Because RAMP1 signaling in KCs appears to be involved in ConA hepatitis, we next isolated KCs and treated them with CGRP. We found that expression of mRNA encoding *Tnf* and *Ifng* in *Ramp1*-deficient KCs stimulated with CGRP (10 nM) was enhanced when compared with that in WT KCs (Fig 6C).

To further examine the interaction between KCs and T cells, we co-cultured isolated KCs (F4/80+ cells) and splenic T cells (CD4+ cells) isolated from WT and *Ramp1^−/−^* mice. Stimulating co-cultured KCs and splenic T cells with ConA further upregulated expression of *Tnf* and *Ifng* when compared with cultured KCs alone or splenic T cells alone (Fig 6D). CGRP reversed the increase in cytokine expression by co-cultured cells from WT mice, but not those from *Ramp1^−/−^* mice (Fig 6D). These results suggest that the interaction between splenic T cells and KCs contributes to ConA hepatitis, which is dependent on RAMP1.

### CGRP suppresses ALT and cytokine production in mice with ConA hepatitis

Finally, to examine whether CGRP suppresses hepatic inflammation elicited by ConA, we investigated the effect of CGRP administration on ALT and cytokine levels. As shown in Fig 7, CGRP administration reduced ALT levels at 24 h; it also reduced expression of mRNA encoding *Tnf* and *Ifng* in the liver and spleen at 1 h.

**Fig 7.**
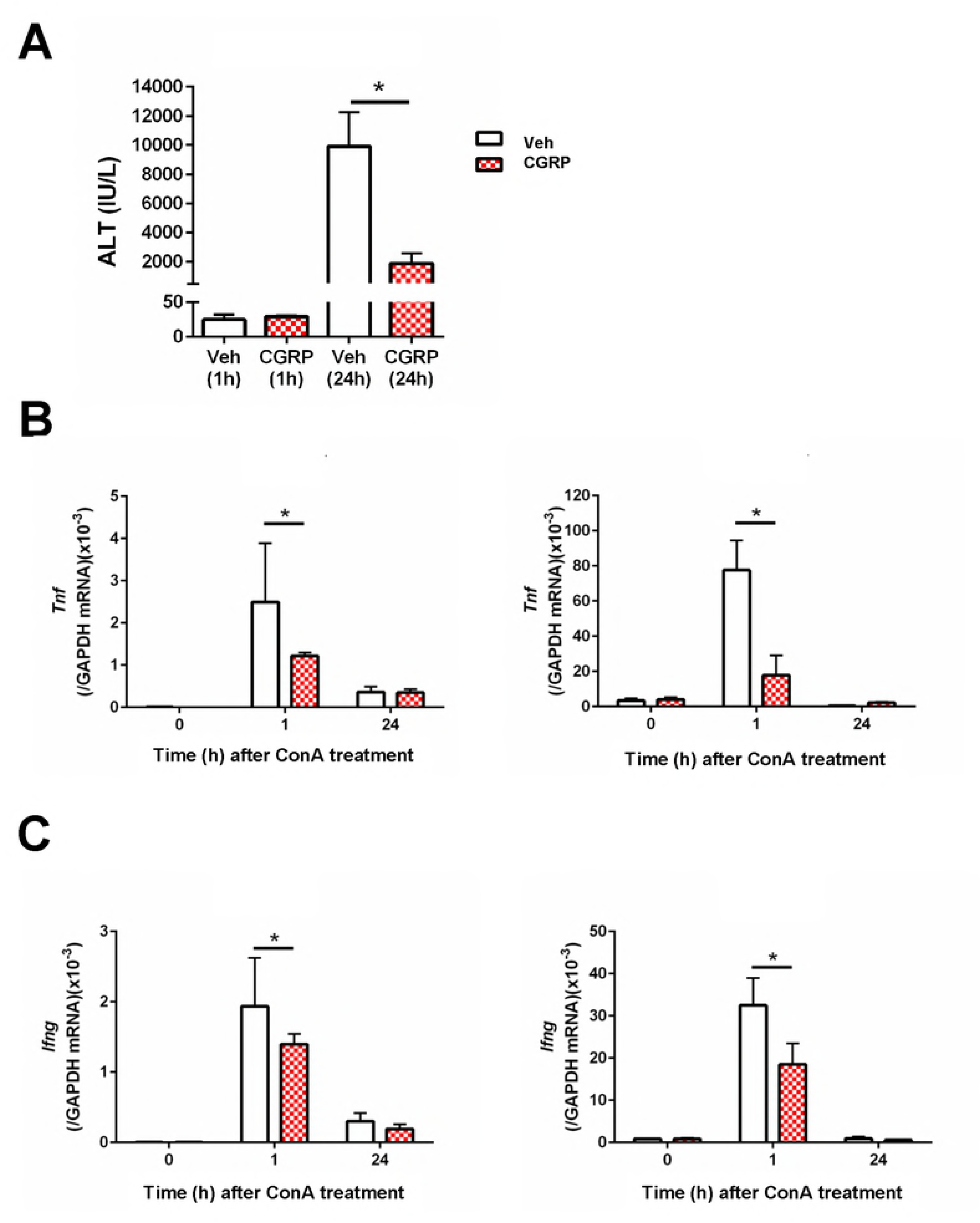
Effects of CGRP treatment on ALT and pro-inflammatory cytokine levels during ConA hepatitis. CGRP or vehicle was administered 30 min before ConA treatment. (a) ALT levels in WT mice treated with CGRP at 1 h and 24 h after ConA administration. Veh, vehicle. Data are expressed as the mean ± SD from 4–6 mice per group. *p < 0.05. (b) Amount of *Tnf* mRNA in the liver and spleen after ConA administration. Data are expressed as the mean ± SD from 4–6 mice per group. *p < 0.05. (c) Amount of Ifng mRNA in liver and spleen after ConA administration. Data are expressed as the mean ± SD from 4–6 mice per group. *p < 0.05.

## Discussion

Sensory neurons communicate actively with the immune system. Here, we demonstrated that RAMP1 signaling plays an immunosuppressive role in ConA-mediated hepatitis by downregulating production of pro-inflammatory cytokines by KCs and splenic T cells. Both T cells and macrophages play key roles in ConA-mediated hepatitis by producing TNFα and IFNγ, both of which cause liver injury. Our data show that splenic T cells trigger hepatic inflammatory responses to ConA and interact with KCs to induce liver injury. KCs are essential for induction and amplification of ConA-mediated hepatitis. RAMP1 signaling inhibits development of ConA-induced hepatitis by inactivating hepatic and splenic immune cells. These findings indicate that RAMP1 is a good therapeutic target for immune-mediated hepatitis.

Although the roles of sensory neuron-immune interactions in response to sterile tissue damage are not fully understood, we showed previously that RAMP1 signaling regulates inflammation by suppressing inflammatory mediators released from immune cells such as dendritic cells, macrophages, and lymphocytes [9,10]. CGRP also contributes to wound healing and secondary lymphedema by affecting RAMP1 signaling in macrophages [21,22]. These findings indicate that CGRP secreted from endings of sensory nerve fibers regulates immune function via RAMP1 signaling.

ConA-induced liver injury is a model for immune-mediated hepatitis, a condition that causes T cell activation followed by increased expression of inflammatory cytokines. During this process, CD4+ T cell-mediated activation of macrophages is a crucial pathogenic parameter [12]. In the liver, endogenous CGRP protects against ConA-induced liver injury [11]. Indeed, *Calca*-deficient (*Calca*^−/−^) mice exhibit aggravation of ConA hepatitis upon reduction of the hepatotrophic cytokine IL-6 and increased apoptosis [11]. However, the role of CGRP receptors in immune-mediated hepatitis is unclear. Here, we demonstrate that RAMP1 attenuates ConA-mediated hepatitis.

CGRP acts on immune cells directly and regulates their production of cytokines in inflammatory lesions [23]. Previously, we reported that RAMP1 is expressed by T cells [10,24], dendritic cells [9], and macrophages [21,22], and that RAMP1 expression by these immune cells is essential for development of inflammation during endotoxemia and colitis. Here, we show that RAMP1 is expressed by hepatic and splenic macrophages and by splenic T cells. These results suggest that immune cells in the liver and spleen are major sites of RAMP1 expression during ConA hepatitis. CGRP and its receptor are expressed in various organs, including liver [15] and spleen [16,17]. In addition, we show that CGRP levels in the liver and spleen fell upon exposure to ConA, and that *Ramp1^−/−^* mice express less CGRP than WT mice. Although CGRP is easily degraded by proteinases present in the circulation immediately after release, maintaining CGRP levels is important for preventing liver injury. Indeed, the data demonstrate that supplementation with CGRP effectively attenuated Con A hepatitis via suppression of pro-inflammatory cytokines.

The spleen is considered a source of pro-inflammatory mediators capable of propagating injury. Accumulating evidence suggests that stimulating the vagus nerve reduces cytokine production by splenic macrophages [25,26]. Binding of the cholinergic receptor (expressed by splenic macrophages) is required for vagus nerve-mediated inhibition of TNFα release [25]. These findings indicate that immune cells and neurons share common signaling molecules, and that the nervous system limits excessive immune activity. Regarding sensory innervation of the spleen, CGRP+ nerve terminals are sited close to the zones in which macrophages and T cells are situated [4]. Therefore, it is important to understand the role of sensory neural innervation of the spleen when examining immune-mediated regulation of tissue injury. The spleen is J * ’ by, or involved in, the pathophysiology of ConA hepatitis. ConA triggers proliferation of and cytokine production by splenic lymphocytes; activated splenic T cells then migrate to the liver to aggravate liver injury [12,27]. The present study shows that splenectomy immediately prior to ConA administration attenuates ConA hepatitis by reducing secretion of pro-inflammatory cytokines. Therefore, the spleen contributes to the early phase of ConA hepatitis by producing pro-inflammatory mediators. However, splenectomy in WT mice did not attenuate ConA hepatitis at 24 h, suggesting that the spleen by itself is not responsible for the severity of the disease. Consistent with this, splenectomy 2 weeks before ConA treatment did not affect liver injury induced by ConA treatment [28]. However, ALT levels in splenectomized *Ramp1^−/−^* mice fell by 50% at 24 h after ConA treatment when compared with those in sham-operated *Ramp1^−/−^* mice, suggesting that RAMP1 signaling in splenocytes improves ConA-mediated hepatitis.

The data also indicate that production of L and M by activated splenic CD4+ T cells in *Ramp1^−/−^* mice was inhibited from as early as 1 h after ConA treatment, at which time no significant liver injury was seen. These results suggest that splenic T cells contribute to ConA hepatitis by producing TNFα and IFNγ, which is dependent on RAMP1 signaling. Alternatively, the spleen (in which T cells are dysregulated after loss of RAMP1-mediated signaling) is the induction site for ConA hepatitis.

We evaluated whether CD4+ T cells are required for ConA hepatitis. Treatment with an anti-CD4 antibody attenuated, but only partly, ConA-mediated hepatitis, indicating that activated CD4+ T cells are responsible for the disease progression. To further determine whether RAMP1 signaling in splenic CD4+ T cells plays a role in development of ConA hepatitis, we examined the effect of adoptively transferring splenic CD4+ T cells from *Ramp1^−/−^* mice. Transfer of Ramp1-deficient splenic CD4+ T cells led to a modest increase in liver injury compared with that in global *Ramp1*-deficient mice. These results suggest that RAMP1 signaling in splenic CD4+ T cells participates in ConA-mediated hepatitis.

In addition to that in splenic T cells, we found that RAMP1 signaling in splenic macrophages suppressed production of L and M- However, splenic macrophages produced fewer pro-inflammatory cytokines than splenic T cells; indeed, flow cytometry analysis revealed that the population of cytokine-secreting cells within splenic macrophages was smaller than that within splenic T cells.

ConA-induced liver injury is initiated not only by specific activation of CD4+ lymphocytes, but also by KCs[13]. Eliminating macrophages markedly attenuates ConA hepatitis [13,14], suggesting that KCs are involved in ConA hepatitis. Consistent with this, we found that deleting macrophages using CL reduced the severity of ConA hepatitis associated with pro-inflammatory mediators in WT and *Ramp1^−/−^* mice. In addition, flow cytometry analysis and *in vitro* studies revealed that RAMP1 signaling downregulated secretion of pro-inflammatory cytokines by KCs. Therefore, production of pro-inflammatory cytokines by KCs is RAMP1-dependent, and RAMP1 signaling in KCs is important for development of ConA hepatitis.

Although macrophage deletion reduced the severity of ConA hepatitis, it also partially suppressed production of pro-inflammatory cytokines. This indicated that cells other than KCs are the source of inflammatory cytokines after ConA stimulation. During the early phase of ConA hepatitis, we detected modest infiltration by hepatic T cells and moderate increases in the number of hepatic CD4+ T cells expressing TNFα and IFNγ in both genotypes, suggesting that RAMP1 signaling in hepatic T cells plays a minor role in early liver injury elicited by ConA. Instead, splenic CD4 T cells, in addition to KCs, were activated as early as 1 h after ConA administration, as evidenced by increased cytokine production, indicating that ConA activates splenic T cells and KCs simultaneously. Furthermore, despite the presence of persistently activated CD4+ T cells in mice treated with CL, the liver was protected from ConA hepatitis, indicating that KCs are required for continuous progression of ConA hepatitis.

After simultaneous activation of splenic T cells and KCs during the course of ConA-mediated hepatitis, migrated splenic T cells induce liver injury by interacting with KCs. This interaction was suggested by results from co-cultures of isolated KCs and splenic T cells. The data showed that expression of pro-inflammatory cytokines in co-culture systems was higher than that in individual cell cultures, and that exacerbated KC and splenic T cell responses to ConA were dependent on RAMP1. Our data also support the notion that TNFα and IFNγ are critical for induction of ConA hepatitis; indeed, we found that antibodies specific for L and M attenuated ConA hepatitis and liver infiltration by inflammatory cells.

Although KCs are crucial for development of ConA hepatitis, their numbers fell significantly at 24 h after ConA treatment. KC-generated cytokines recruited Ly6C- or CD68-positive monocytes/macrophages, leading to extensive liver injury during the late phase of ConA hepatitis. Of interest, the liver of WT mice was infiltrated by Ly6C^high^/CD11b^high^/F4/80^high^ cells, while that of *Ramp1^−/−^* mice displayed accumulation of Ly6C^high^/CD11b^high^/F4/80^low^ cells. This suggests that susceptibility of *Ramp1^−/−^* mice to ConA hepatitis is related to alterations in macrophage phenotype. Preliminary studies showed that the survival rate of WT mice with ConA hepatitis was 100%, and that ALT levels at 48 h were 206 ± 40 IU/L (n = 6). By contrast, survival rates for *Ramp1^−/−^* mice were significantly lower. Considering the different mortality rates and mechanisms of liver injury between the two genotypes, it may be that Ly6C^high^/CD11b^high^/F4/80^high^ monocyte-derived macrophages in WT mice contribute to resolution of inflammation after ConA hepatitis, while Ly6C^high^/CD11b^high^/F4/80^low^ monocyte-derived macrophages in *Ramp1^−/−^* mice exacerbate inflammation in response to ConA. Indeed, Ly6C^high^/CD11b^high^/F4/80^high^ monocyte-derived macrophages are involved in liver repair after acetaminophen hepatotoxicity [29]. In addition, we recently reported that Ly6C^high^/CD11b^high^/F4/80^low^ cells recruited to the liver display a pro-inflammatory macrophage phenotype and delay liver repair after hepatic ischemia/reperfusion [20]. The phenotypes of macrophages recruited to injured livers at 24 h post-ConA treatment were different in WT and *Ramp1^−/−^* mice, indicating that RAMP1 signaling is involved in macrophage polarization. Further studies are needed to elucidate the role of RAMP1 signaling in resolution of liver inflammation and liver repair during ConA hepatitis.

In conclusion, the present study demonstrates that RAMP1 signaling in KCs plays a crucial role in preventing and/or limiting immune-mediated hepatitis; this may have important implications for the treatment of patients with autoimmune liver inflammation. Considering that dysregulation of RAMP1 function has marked effects on the outcome of murine liver inflammation, selective agonists of RAMP1 might be a therapeutic option for immune-mediated hepatitis. Therefore, restorative therapies aiming at restoring RAMP1 activity in the immune cells might modulate liver injury.

## Acknowledgements

We thank Michiko Ogino and Kyoko Yoshikawa for their technical assistance.

